# Pre-attentive processing of neutral and emotional sounds in congenital amusia

**DOI:** 10.1101/2020.08.05.238204

**Authors:** Agathe Pralus, Marie Gomot, Jackson Graves, Fanny Cholvy, Lesly Fornoni, Barbara Tillmann, Anne Caclin

## Abstract

Congenital amusia is a life-long deficit of musical processing. This deficit can extend to the processing of language and in particular, emotional prosody. In a previous behavioral study, we revealed that while amusic individuals had difficulties in explicitly recognizing emotions for short vowels, they rated the emotional intensity of these same vowels as did their matched control participants. This finding led to the hypothesis that congenital amusics might be impaired for explicit emotional prosody recognition, but not for its implicit processing. With the aim to investigate amusics’ automatic processing of prosody, the present study measured electroencephalography (EEG) when participants listened passively to vowels presented within an oddball paradigm. Emotionally neutral vowel served as the standard and either emotional (anger and sadness) or neutral vowels as deviants. Evoked potentials were compared between participants with congenital amusia and control participants matched in age, education, and musical training. The MMN was rather preserved for all deviants in amusia, whereas an earlier negative component was found decreased in amplitude in amusics compared to controls for the neutral and sadness deviants. For the most salient deviant (anger), the P3a was decreased in amplitude for amusics compared to controls. These results showed some preserved automatic detection of emotional deviance in amusia despite an early deficit to process subtle acoustic changes. In addition, the automatic attentional shift in response to salient deviants at later processing stages was reduced in amusics in comparison to the controls. In the three ERPs related to the deviance, between-group differences were larger over bilateral prefrontal areas, previously shown to display functional impairments in congenital amusia. Our present study thus provides further understanding of the dichotomy between implicit and explicit processing in congenital amusia, in particular for vocal stimuli with emotional content.

## Introduction

Congenital amusia, also known as tone-deafness, is a life-long deficit of music processing. This deficit seems to affect one to two percent of the general population (Peretz et al., 2007; Peretz & Vuvan, 2017), with potentially genetic origins (Peretz et al., 2007). Individuals with congenital amusia show no hearing impairments or brain lesions that could explain their deficit. They are usually unable to sing in tune or detect an out-of-key note (see Peretz, 2016; Tillmann et al., 2015 for reviews). Several studies have revealed a specific pitch processing deficit in congenital amusia, with pitch perception tasks (Hyde & Peretz, 2004; Peretz et al., 2003) and pitch memory tasks (Albouy et al., 2016; Graves et al., 2019; Tillmann, Lévêque, et al., 2016; Williamson & Stewart, 2010). The pitch deficit was observed for non-musical material, such as isolated pitches or tone pairs (Albouy et al., 2016; Foxton et al., 2004; Peretz et al., 2009), as well as tone sequences or melodies (see Tillmann et al., 2015 for a review). As pitch processing is relevant beyond the musical domain, the investigation of congenital amusia has been extended to speech perception abilities. While some early studies did not report any deficit of speech processing in amusic individuals (Ayotte et al., 2002; Tillmann et al., 2009; Williamson & Stewart, 2010), more recent studies have revealed specific impairments of speech contour perception and intonation recognition in congenital amusia (Jiang et al., 2010; Liu et al., 2015, 2017; Nan et al., 2016; Nguyen et al., 2009; Tillmann, Burnham, et al., 2011; Tillmann, Rusconi, et al., 2011).

As pitch is essential to process emotions both in speech and music, some studies have started to investigate emotional processing in congenital amusia. Regarding musical emotion perception, congenital amusics have demonstrated either a mild impairment or no impairment in recognition tasks (Gosselin et al., 2015; Lévêque et al., 2018; Marin et al., 2015), but have shown preserved intensity ratings of the emotions (Lévêque et al., 2018). Regarding emotion perception in speech, referred to as emotional prosody, congenital amusics have demonstrated a mild deficit of recognition (Lima et al., 2016; Pralus et al., 2019; Thompson et al., 2012), which was more pronounced for short vowels (with few acoustic cues) than long sentences (Lolli et al., 2015; Pralus et al., 2019). This recognition deficit was the largest for sadness stimuli (Pralus et al., 2019; Thompson et al., 2012), which tended to be confounded with neutral stimuli (Pralus et al., 2019). Interestingly, when congenital amusics were asked to rate the intensity of emotional prosody stimuli, they did not show any deficit compared to matched controls, even for vowels (Pralus et al., 2019). Intensity ratings of emotions can be given without precise categorical representation of the emotion or explicit labeling, suggesting some preserved implicit processing of emotions in amusia (Lévêque et al., 2018; Pralus et al., 2019). For music material, preserved implicit processing of pitch in amusia has been reported, even though explicit processing has been shown to be disrupted (Lévêque et al., 2018; Omigie et al., 2013; Pralus et al., 2019; Tillmann et al., 2012, 2014; Tillmann, Lalitte, et al., 2016). For instance, congenital amusics were able to process pitch changes as well as pitch incongruity (Peretz et al., 2009; Zendel et al., 2015), even though they were unable to detect these changes or incongruities when explicitly asked to do so (Moreau et al., 2009; Omigie et al., 2012; Tillmann, Lévêque, et al., 2016). This recent research suggests congenital amusia to be a disorder of consciousness related to pitch representations (Albouy et al., 2016; Marin et al., 2015; Moreau et al., 2009, 2013; Omigie et al., 2013; Peretz, 2016; Peretz et al., 2009; Stewart, 2011; Tillmann, Lalitte, et al., 2016).

Aiming to further investigate this hypothesis, in the present study, we recorded several electrophysiological measures that have been previously used to explore implicit processes in the typical and the pathological brain. One target measure, which reflects pre-attentional processes in the brain, is the well-studied Mismatch Negativity (MMN) (Näätänen et al., 2007; Näätänen & Alho, 1995). This negative ERP component emerges when a deviant event appears in a repetitive auditory sequence (referred to as the oddball paradigm). It is considered to be automatic as it can be recorded even when participants are actively engaged in another task (Näätänen et al., 2007; Näätänen & Alho, 1995). The MMN signal is very robust, stable, and found in most control participants at the individual level (Chen et al., 2018; Kraus et al., 1992). It is generally reported to originate from supratemporal and frontal cortical regions (Näätänen et al., 2007; Näätänen & Alho, 1995). An MMN can be induced not only by simple acoustic deviants, as classically studied (Näätänen et al., 2007; Peretz et al., 2005), but also by emotional deviant events (Goydke et al., 2004). For emotional prosodic material, such as vowels, an MMN can be induced by an emotional deviant, compared to a neutral standard (Carminati et al., 2018; Charpentier et al., 2018). This emotional MMN occurs generally at a shorter latency and is larger than for neutral deviant (Schirmer et al., 2005, 2016).

For non-emotional material, amusics’ automatic brain response to acoustic changes has been studied in passive listening paradigms with pitch tone deviants or tone-language stimuli (Fakche et al., 2018; Moreau et al., 2009, 2013; Nan et al., 2016; Omigie et al., 2013; Zhang & Shao, 2018). Using pitch change passive paradigms, amusics’ early change-related evoked potentials, such as the MMN, were decreased in amplitude in comparison to controls for small pitch changes (Fakche et al., 2018; Moreau et al., 2009, 2013). When the pitch change was large enough (200 cents), the MMN seemed to be preserved in amusics (Moreau et al., 2009, 2013). Omigie et al. (2013) used real melodies to investigate amusics’ and controls’ brain responses as a function of the degree of expectedness of the notes (Omigie et al., 2013). The results revealed that with increased unexpectedness the early negativity (in the N1 latency range) increased for controls, but not for amusics. It suggests a deficit in the processing of musical structures at early processing stages, in keeping with the results of Albouy et al. (2013) in an active short-term memory task for melodies. When tone-language stimuli were used, amusics did not demonstrate any decrease of MMN in response to lexical tones (Nan et al., 2016; Zhang & Shao, 2018).

In active paradigms, similar result patterns regarding early brain response to different acoustic changes in congenital amusia have been observed (Braun et al., 2008; Peretz et al., 2005, 2009). For pitch change detection tasks, the MMN was decreased for amusics (compared to controls) only for small pitch deviants (Peretz et al., 2005, 2009). The replacement of the correct tone with an incorrect (out-of-tune) deviant tone at the end of familiar melodies revealed a decreased early negativity in amusics compared to controls (Braun et al., 2008). For language material, in particular intonation processing with statements and questions, the early negativity was preserved in amusics, but the N2 response was decreased in response to incongruent pairs of tones (Lu et al., 2015).

Some alterations in deviance detection have been observed also for later components, such as the P3 (Braun et al., 2008; Lu et al., 2015; Moreau et al., 2009, 2013; Peretz et al., 2009; Zhang & Shao, 2018). For pitch change detection tasks using tones, a decreased P3 was observed for amusics (in comparisons to controls) only for small pitch changes (25 cents), but not otherwise (Braun et al., 2008; Moreau et al., 2013). For lexical tone changes, smaller P3a and P3b were observed in amusics compared to controls for small lexical tone changes (high rising vs. low rising tone) (Zhang & Shao, 2018).

Overall, some results have shown decreased early electrophysiological markers related to pitch deviance detection in congenital amusia, mostly for small pitch changes, and sometime together with a reduction of the subsequent P3a. However, the pattern of automatic pitch processing in speech and music in congenital amusia still needs further investigation.

Our previous behavioral study investigating emotional prosody in congenital amusia has suggested preserved implicit prosody processing (Pralus et al., 2019). With the aim to further investigate amusics’ automatic processing of prosody, the present study measured electroencephalography (EEG) when participants listened passively to vowels presented within an oddball paradigm. Emotionally neutral vowel served as the standard and either emotional (anger and sadness) or neutral vowels as deviants. Evoked potentials were compared between participants with congenital amusia and control participants matched in age, education, and musical training. Emotional deviants (anger and sadness) and neutral deviant were chosen from the material of our previous study (Pralus et al., 2019) aiming for similar F0 difference compared to the neutral standard. Anger was the best recognized emotion by amusics whereas sadness was not well recognized and often confused with neutrality. Hence, these two emotional deviants had different patterns of recognition in the two participant groups, while intensity ratings were similar across groups for these stimuli. We hypothesized that early automatic processing of emotion deviancy will be impaired in amusics compared to controls, with potentially different responses to neutral and emotional deviancy in these two groups.

## Material and Methods

### Participants

Nineteen amusic participants and twenty-one control participants matched for gender, age, laterality, education, and musical training (as defined by years of instruction of an instrument) at the group level were included in the study (**Table 1**). They all gave written informed consent to participate in the experiment. Prior to the main experiment, all participants were tested with a subjective audiometry, the Montreal Battery of Evaluation of Amusia (Peretz et al., 2003) to diagnose amusia, and a Pitch Discrimination Threshold (PDT) test (Tillmann et al., 2009). A participant was considered amusic if he/she had a global MBEA score below 23 (maximum score = 30) and/or a MBEA pitch score (average of the first three subtests of the MBEA) inferior to 22 (maximum score = 30). All control participants had a global MBEA score above 24.5 and a MBEA pitch score above 23.3 (see **Table 1**). All participants had normal hearing (hearing loss inferior to 30 dB at any frequency in both ears). Study procedures were approved by a national ethics committee. Participants provided written informed consent prior to the experiment and were paid for their participation.

**Table 1:** Characteristics of the participants in both groups. The MBEA (Montreal Battery for the Evaluation of Amusia, Peretz et al., 2003) score corresponds to the average of the six subtests of the battery (maximum score = 30, cut off: 23). Pitch mean score corresponds to the average of the three pitch subtests in the MBEA (scale, contour and interval, cut off: 22). Note that a participant was considered as amusic if any of these two measures (MBEA score, MBEA pitch score) was below the cut-off. PDT: Pitch Discrimination Threshold (see Tillmann et al., 2009). For each variable (except sex and handedness), the mean value in each group is reported along with the standard deviation in parentheses. Groups were compared with t.tests (two sided), except for sex and handedness where a Chi2 test was used (Qobs=0.35 and Qobs=0.3, respectively)

|  | **Amusics (n=19)** | **Controls (n=21)** | **p-value (group comparison)** |
| --- | --- | --- | --- |
| **Age (years)** | 30.7 ($\pm 14.38$) | 32.33 ($\pm 14.5$) | 0.72 |
|  | Min: 18 | Min: 19 |  |
|  | Max: 56 | Max: 64 |  |
| **Education (years)** | 15 ($\pm 2.67$) | 15.23 ($\pm 2.19$) | 0.76 |
|  | Min: 10 | Min: 12 |  |
|  | Max: 20 | Max: 20 |  |
| **Musical training (years)** | 0 | 0.048 ($\pm 0.22$) | 0.33 |
|  |  | Min: 0 |  |
|  |  | Max: 1 |  |
| **Sex** | 9M 10F | 8M 13F | 0.55 |
| **Handedness** | 5L 14R | 4L 17R | 0.58 |
| **MBEA score** | 22.02 ($\pm 1.8$) | 26.45 ($\pm 1.04$) | **<0.001** |
|  | Min: 16.83 | Min: 24.8 |  |
|  | Max: 24.5 | Max: 28.5 |  |
| **MBEA pitch score** | 21.05 ($\pm 1.97$) | 26.6 ($\pm 1.42$) | **<0.001** |
|  | Min: 15.67 | Min: 23.33 |  |
|  | Max: 23.67 | Max: 28.67 |  |
| **PDT (semitones)** | 1.33 ($\pm 1.48$) | 0.29 ($\pm 0.15$) | **0.007** |
|  | Min: 0.11 | Min: 0.08 |  |
|  | Max: 4.99 | Max: 0.71 |  |

### Stimuli

Four vowels /a/ were selected from a larger material set, all produced with female voices (Charpentier et al., 2018), and used in a previous behavioral study with amusic (N=18) and control (N=18) participants (Pralus et al., 2019). All stimuli lasted 400 ms and were equalized in RMS amplitude. The stimuli were selected based on their recognition scores in the behavioral task (Pralus et al., 2019, see **Table 2**) as follows: the neutral deviant and standard were equally well recognized by all participants; the anger deviant was selected as an easy deviant (equally well-recognized by both groups); the sadness deviant was selected as a difficult deviant for amusics. We added the constraint that all stimuli should be similar in pitch and should have received similar intensity ratings (for emotional stimuli) (see **Table 2** for details). Acoustic parameters (pitch mean, spectral flux mean, brightness mean, roughness mean, inharmonicity mean, and attack time) of the stimuli were computed with the MIR toolbox (Lartillot & Toiviainen, 2007); **Table 2**). Each parameter (except Attack Time) was computed with a temporal frame of 50ms by default. We then computed the average of each parameter across time (see **Table 2**).

**Table 2:** Acoustic parameters of the stimuli and associated behavioral data from Pralus et al., *(2019)*. The acoustic parameters were computed with the MIR Toolbox (Lartillot & Toiviainen, 2007), with a temporal frame of 50ms. a.u.: arbitrary units. Percentage of correct emotion recognition and intensity ratings (on a scale from 1 to 5) for these stimuli are from Pralus et al. *(2019)* and were obtained from 18 congenital amusics and 18 matched controls. NA: not applicable, no intensity ratings was given for neutral stimuli.

| *Acoustic parameters* | **Neutral standard** | **Neutral deviant** | **Sadness deviant** | **Anger deviant** |
| --- | --- | --- | --- | --- |
| *Pitch mean (Hz)* | 241 | 199 | 228 | 278 |
| *Spectral flux mean (a.u.)* | 17.19 | 9.33 | 25.75 | 68.70 |
| *Brightness mean (a.u.)* | 0.20 | 0.13 | 0.23 | 0.27 |
| *Roughness mean (a.u.)* | 22.98 | 38.09 | 14.21 | 114.80 |
| *Inharmonicity mean (a.u.)* | 0.18 | 0.17 | 0.27 | 0.45 |
| *Attack time (s)* | 0.028 | 0.039 | 0.056 | 0.13 |
| ***Behavioral data***<br>***(Pralus et al., 2019)*** |  |  |  |  |
| *% Correct recognition in Controls* | 100 | 83 | 94 | 72 |
| *% Correct recognition in Amusics* | 94 | 83 | 56 | 67 |
| *Mean Intensity ratings in Controls* | NA | NA | 2.8 | 2.6 |
| *Mean Intensity ratings in Amusics* | NA | NA | 2.8 | 2.4 |

### Procedure

The experiment took place in a sound-attenuated room. Participants watched a silent movie with subtitles, they were told to not pay attention to the sounds played over headphones. The recording session lasted 45 minutes.

### EEG recordings and ERP measurements

The entire experimental paradigm was composed of three oddball blocks, each with one type of deviant (Neutral, Sadness, Anger) and one block with equiprobable stimuli. For each oddball block, 700 standards and 140 deviants were played. Two consecutive deviants were separated by at least three standards. During the equiprobable block, each of the 4 stimuli were played equally often (144 times each, 576 stimuli in total), with no more than two repetitions of the same stimulus in a row. The stimulus onset asynchrony (SOA) was always 700 ms.

EEG was recorded using 31 active electrodes (BrainAmp/Acticap, Brain Products, Germany) with a nose reference, with a sampling frequency of 1000Hz (bandwidth 0.016-1000 Hz). Eye movements were recorded with an electrode under the left eye (offline re-referenced to Fp1). ELAN software was used for EEG signal processing (Aguera et al., 2011). Band-stop filters centered around 50Hz and 150Hz were applied to the EEG signal to remove power line artifacts. Independent Component Analysis was performed on the EEG signal to remove artifacts due to eye movements and heartbeat (Delorme & Makeig, 2004). Averaging was done for each deviant and standard separately, in the three oddball blocks and the equiprobable block. Standards occurring after a deviant were not averaged. Averaging was done on a 700ms time-window (from −200 ms to 500 ms around stimulus onset). Trials with peak-to-peak amplitude variation exceeding 150 µV at any electrode were rejected. Noisy electrodes were interpolated. A 2-30Hz band-pass Butterworth filter (order 4) was applied to the evoked potentials. ERPs were baseline-corrected by subtracting the average of the signal in the 100ms before the stimulus. The difference wave for each type of deviant (Neutral, Sadness, Anger) was obtained by subtracting the response to the deviant from the response to the standard in the same block of the oddball paradigm^1^. Grand-averaged curves were obtained for both groups (Amusics and Controls). The emergence of deviance-related ERPs (MMN and P3a in particular) was assessed with the comparison of deviant and standard ERPs using a nonparametric cluster-based permutation analysis (1000 permutations), in each group, for each of the three deviants. A first threshold of p<0.05 was used for permutation-based paired t-tests for each sample. Clusters were labeled as significant for p<0.05 at the end of the permutations, controlling for multiple comparisons in space (31 electrodes) and time. Based on the union of these emergence tests in both groups, two or three time windows of interest were selected for each emotion. For neutral deviant, three time-windows were selected: 67-130ms, 130-205ms, 225-310ms. For sadness deviant, three time-windows were selected: 77-140ms, 140-200ms, 220-295ms. For anger deviant, two time-windows were selected: 113-205ms, 217-299ms. The first window corresponds to an early negativity at the latency of the N1 (neutral and sadness deviant only), the next one to the MMN, and the last to the P3a.

### Statistical analysis

Based on the emergence tests described above, a set of fronto-central electrodes was selected for the main analysis. Average amplitude for electrode sites along the antero-posterior axis (four levels) and for the two sides (pre-frontal=Fp1, Fp2, frontal=F3, F4, fronto-central=FC1, FC2, central=C3, C4, odd numbers correspond to electrodes on the left side, even numbers on the right side) were computed for each participant, for each type of deviant, in each of the time windows of interest. For each emotion (neutral, sadness, anger) and for each time-window (early negativity, MMN, P3, except for anger for which there was no early negativity), a Bayesian repeated-measures Analysis of Variance (ANOVA) was performed with group (Amusics, Controls) as a between-subjects factor, and localization (Fp, F, FC and C) and side (left, right) as within-subject factors^2^.

We report Bayes Factor (BF) as a relative measure of evidence. To interpret the strength of evidence (according to Lee & Wagenmakers, 2014), we considered a BF under three as weak evidence, a BF between three and 10 as positive evidence, a BF between 10 and 100 as strong evidence and a BF higher than 100 as a decisive evidence. BF_10_ indicates the evidence of H1 (a given model) compared to H0 (the null model), and BF_inclusion_ indicates the evidence of one effect over all models. As no post-hoc tests with correction for multiple comparison have as yet been developed for Bayesian statistics (Wagenmakers et al., 2017, 2018), we used t-tests with Holm-Bonferroni correction for multiple comparisons.

### Data availability

Raw data were generated at Lyon Neuroscience Research Center (France). Derived data supporting the findings of this study are available from the corresponding author upon request.

## Results

Based on the emergence tests, three deviance-related ERPs were identified in the difference curves (**Figures 1–3**): (1) an early negativity was observed, namely a negative fronto-central deflection in a time-window of ~70-140ms after the stimulus onset; (2) the MMN was identified as the negative fronto-central deflection in a time-window of ~140-200ms after the stimulus onset, associated with the typical polarity inversion at the mastoids; (3) the P3a was identified as the positive fronto-central deflection in a time-window of ~220-300ms after the stimulus onset, with a polarity inversion at the mastoids. See Figures 1–3A for averaged curves over fronto-central sites and **Figures 1–3B** for topographies. For the Anger deviant difference curve, only two emergence windows were retrieved, corresponding to the MMN and the P3a. For precise emergence windows for each emotion and each group, see **Figures 1–3C**. Overall, the morphology of deviance-related responses was slightly different across emotions. In particular, there were differences in the latencies of the ERPs across emotions, these latencies were similar between groups.

**Figure 1:**
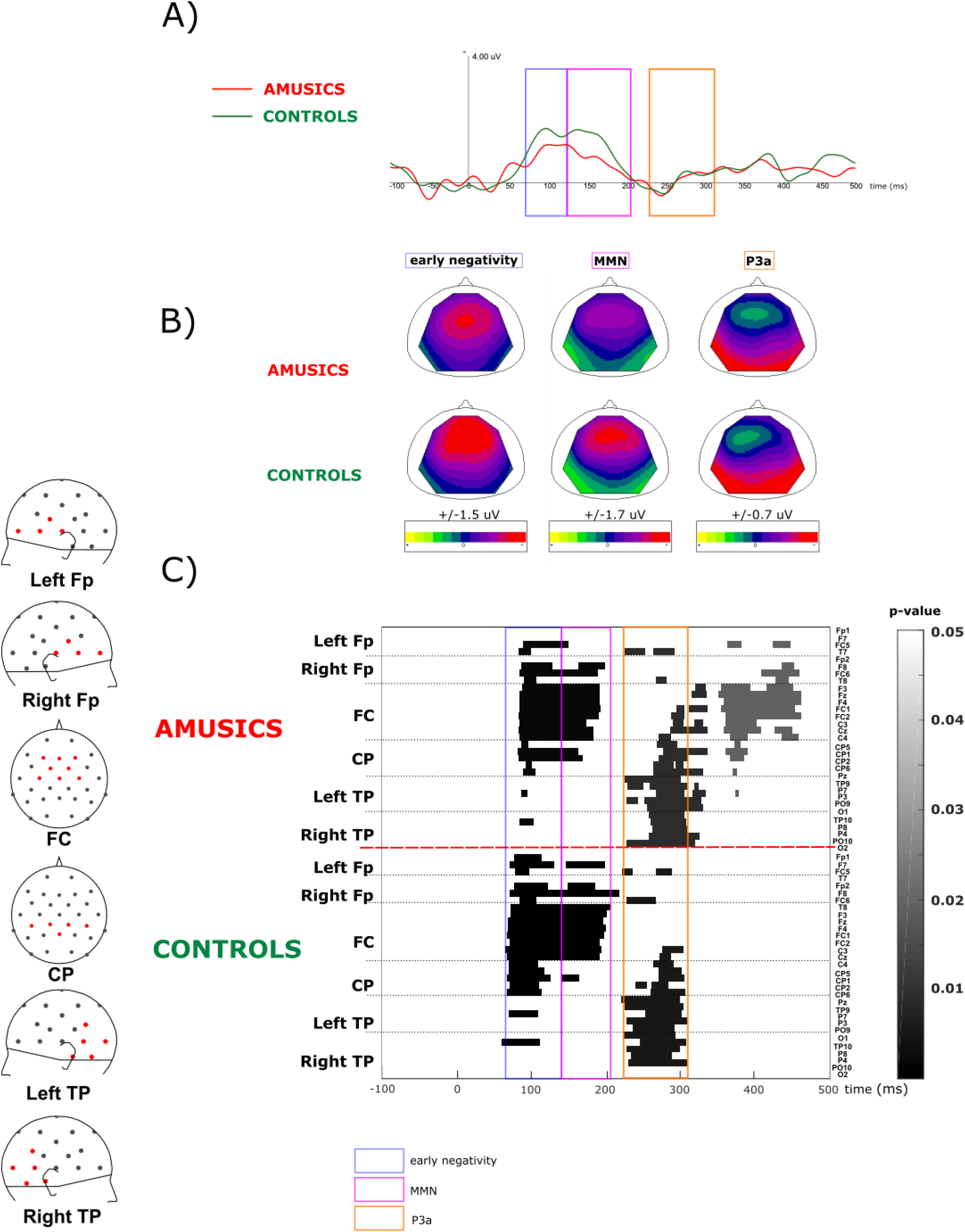
Evoked response to an emotionally neutral deviant in Amusics and Controls. **A)** Average curve of eight fronto-central electrodes (Fp1, Fp2, F3, F4, FC1, FC2, C3, C4) of the response to the neutral deviant minus the response to the neutral standard, for amusics and controls, negativity is up. **B)** Topographies for the three evoked potentials (early negativity, MMN and P3) over the emergence windows identified below, separately for amusics and controls. Amplitude scale is indicated for each ERP. **C)** Emergence of evoked responses for amusics and controls for each electrode, grouped by topography, emergence windows used for the analysis are in blue for early negativity (67-130ms), pink for the MMN (130-205ms), orange for the P3a (225-310ms). Fp= pre-frontal, FC=fronto-central, CP=centro-parietal, TP=temporo-parietal.

Only averaged curves with neutral and sadness deviants showed an early negativity on fronto-central electrodes (**Figures 1–2B**), which is at the latency of the N1: For the neutral deviant, the emergence was between 67 and 130ms, whereas it was later for the sadness deviant, between 77 and 140ms. As habituation of the N1 was visible on standards in the oddball blocks (**Figure S2**), this early negativity might mainly reflect the different degrees of habituation of the N1 between standards and deviants. This effect was possibly less pronounced for the anger deviant, which had a slower attack time (see **Table 2**), and resulted in later auditory ERPs (see in particular the delay in the P50 with respect to the other stimuli in the equiprobable block, **Figure S1**). At the MMN latency, a typical ERP was observed for the three types of deviants, emerging at different latencies for each emotion: 130-205ms for neutrality, 140-200ms for sadness, 113-205ms for anger. For anger, the MMN peak was larger than the other two for both groups. At the P3a latency, the three deviance-related ERPs had an emerging peak with different latencies for each emotion: 225-310ms for neutrality, 220-295 for sadness, 217-299ms for anger.

Based on these observations, and in particular, that ERPs and their latencies were not identical in the three emotions, the main analyses were performed separately by emotion and by component.

### Response to a neutral deviant (Figure 1)

#### 1. Early negativity

After comparison to the null model, the best model showing decisive evidence was the model with the main effects of Localization, Group, and the interaction between the two (BF10=1.2e+9). This model was 2.3 times better than the model with the main effect of Localization (BF10=5.3e+8), 3.6 times better than the model with the main effects of Localization and Group (BF10=3.37e+8), and 4.9 times better than the model with the main effects of Localization, Group, Side and the interaction between Group and Localization (BF10=2.44e+8). The best model was at least 11.5 times better than the other models (BF10<1.04e+8). This was confirmed by a decisive specific effect of Localization (BFinclusion=4.8e+8), a positive effect of the interaction between Localization and Group (BFinclusion=3.06) and no other specific effects (BFinclusion<1.07). According to post-hoc tests, amplitude at Fp sites was smaller than amplitudes at F, FC and C (all pcorr<0.025), amplitude at FC was higher than amplitudes at F and C (all pcorr<0.025). Amusics had a significantly smaller early negativity than controls. Specifically, amusics had smaller amplitude at Fp compared to amplitudes at F, FC, C (all pcorr<0.001), whereas no such pattern was observed in controls (all pcorr>0.39).

#### 2. MMN

After comparison to the null model, the best model showing decisive evidence was the model with the main effect of Localization (BF10=3.33e+14). This model was 1.3 times better than the model with the main effects of Localization and Group (BF10=2.52e+14), 7.1 times better than the model with the main effects of Localization and Side (BF10=4.66e+13), and 9.3 times better than the model with the main effects of Localization, Group, and Side (BF10=3.59e+13). The best model was at least 12 times better than the other models (BF10<2.75e+13). This was confirmed by a decisive specific effect of Localization (BFinclusion=6.43e+13), and no other specific effects (BFinclusion<0.31). According to post-hoc tests, amplitude at Fp and C sites was smaller than amplitudes at F, FC (all pcorr<0.001). The Group effect emerging in the second best model showed that amusics tended to have a smaller MMN than controls.

#### 3. P3a

After comparison to the null model, the best model showing decisive evidence was the model with the main effects of Localization and Side (BF10=2.05e+10). This model was 1.9 times better than the model with the main effects of Localization, Side, and Group (BF10=1.07e+10), and 2.3 times better than the model with the main effects of Localization, Side, Group, and the interaction between Side and Group (BF10=8.86e+9). The best model was at least 10 times better than the other models (BF10<2.05e+9). This was confirmed by a decisive specific effect of Localization (BFinclusion=1.57e+9), a positive effect of Side (BFinclusion=6.01), and no other specific effects (BFinclusion<0.59). Amplitudes were larger over left side than right side. The Group effect emerging in the second best model showed that amusics tended to have a bigger P3a than controls. According to post-hoc tests, amplitudes at Fp and C sites were smaller than amplitudes at F and FC (all pcorr<0.001).

### Response to a sadness deviant (Figure 2)

#### 1. Early negativity

After comparison to the null model, the best model showing decisive evidence was the model with the main effects of Localization, Group and the interaction between the two (BF10=1.74e+9). This model was only 1.04 times better than the model with the main effect of Localization (BF10=1.68e+9), and 1.43 times better than the model with the main effects of Localization and Group (BF10=1.22e+9). The best model was at least 8.3 times better than the other models (BF10<2.1e+8). This was confirmed by a decisive specific effect of Localization (BFinclusion=9.36e+8), and no other specific effects (BFinclusion<1.37). Amusics had a smaller early negativity than controls. According to post-hoc tests, amplitude at Fp sites was smaller than amplitudes at F, FC (both pcorr<0.001), amplitudes at F and C were smaller than amplitude at FC (both pcorr<0.007). Specifically, amusics had smaller amplitude at Fp compared to amplitudes at F, FC, C (all pcorr<0.004), whereas controls had smaller amplitude at Fp compared only to FC (pcorr=0.021).

**Figure 2:**
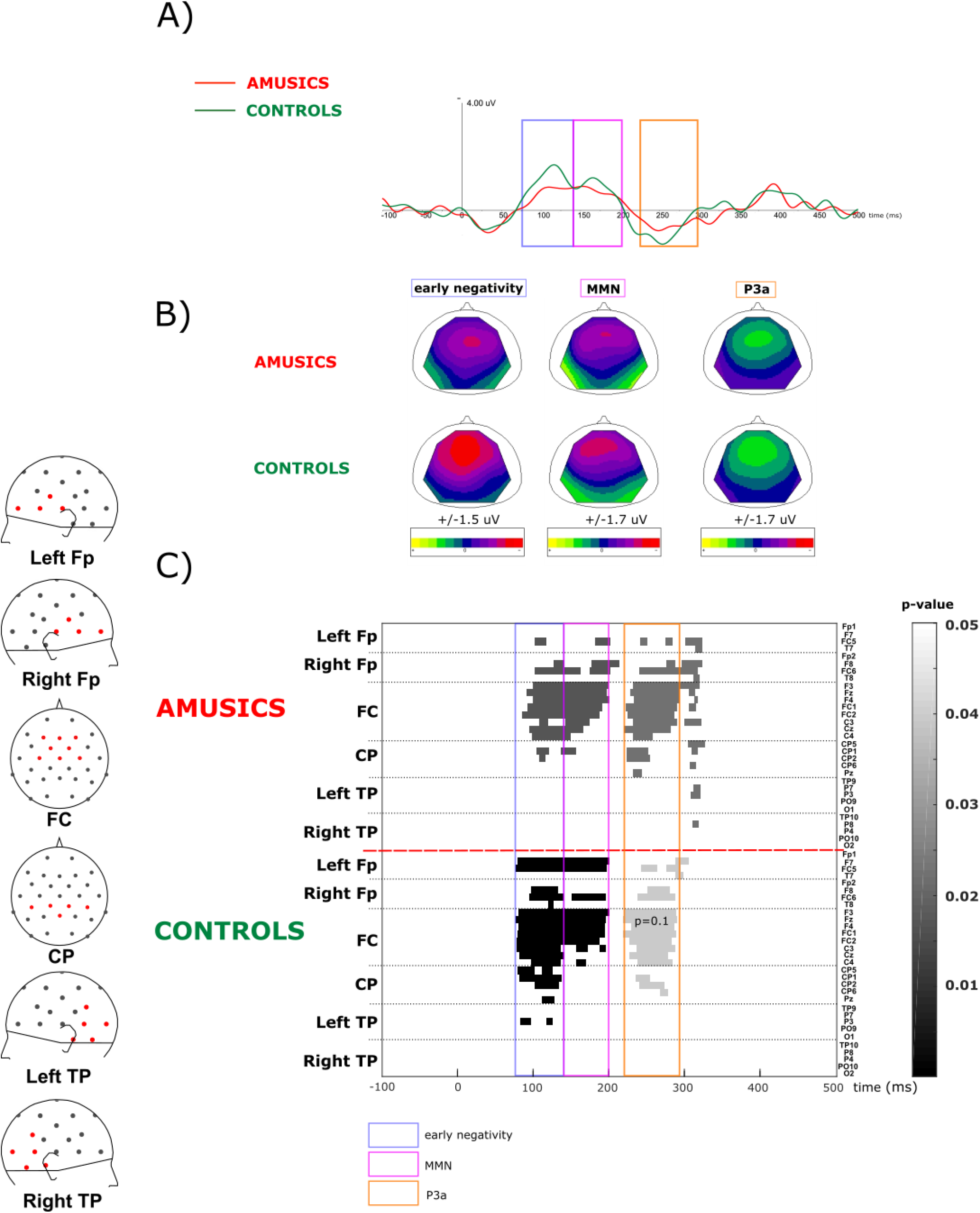
Evoked response to an emotional sadness deviant in Amusics and Controls. **A)** Average curve of eight fronto-central electrodes (Fp1, Fp2, F3, F4, FC1, FC2, C3, C4) of the response to the sadness deviant minus the response to the neutral standard, for amusics and controls, negativity is up. **B)** Topographies for the three evoked potentials (early negativity, MMN and P3) over the emergence windows identified below, separately for amusics and controls. Amplitude scale is indicated for each ERP. **C)** Emergence of evoked responses for amusics and controls for each electrode, grouped by topography, emergence windows used for the analysis are in blue for early negativity (77-140ms), pink for the MMN (140-200ms), orange for the P3a (220-295ms). The P3a only emerged in the amusic group at a pvalue of 0.05 for the permutation test, if the pvalue was set at 0.1 it also emerged in the control group. Fp= pre-frontal, FC=fronto-central, CP=centro-parietal, TP=temporo-parietal.

**Figure 3:**
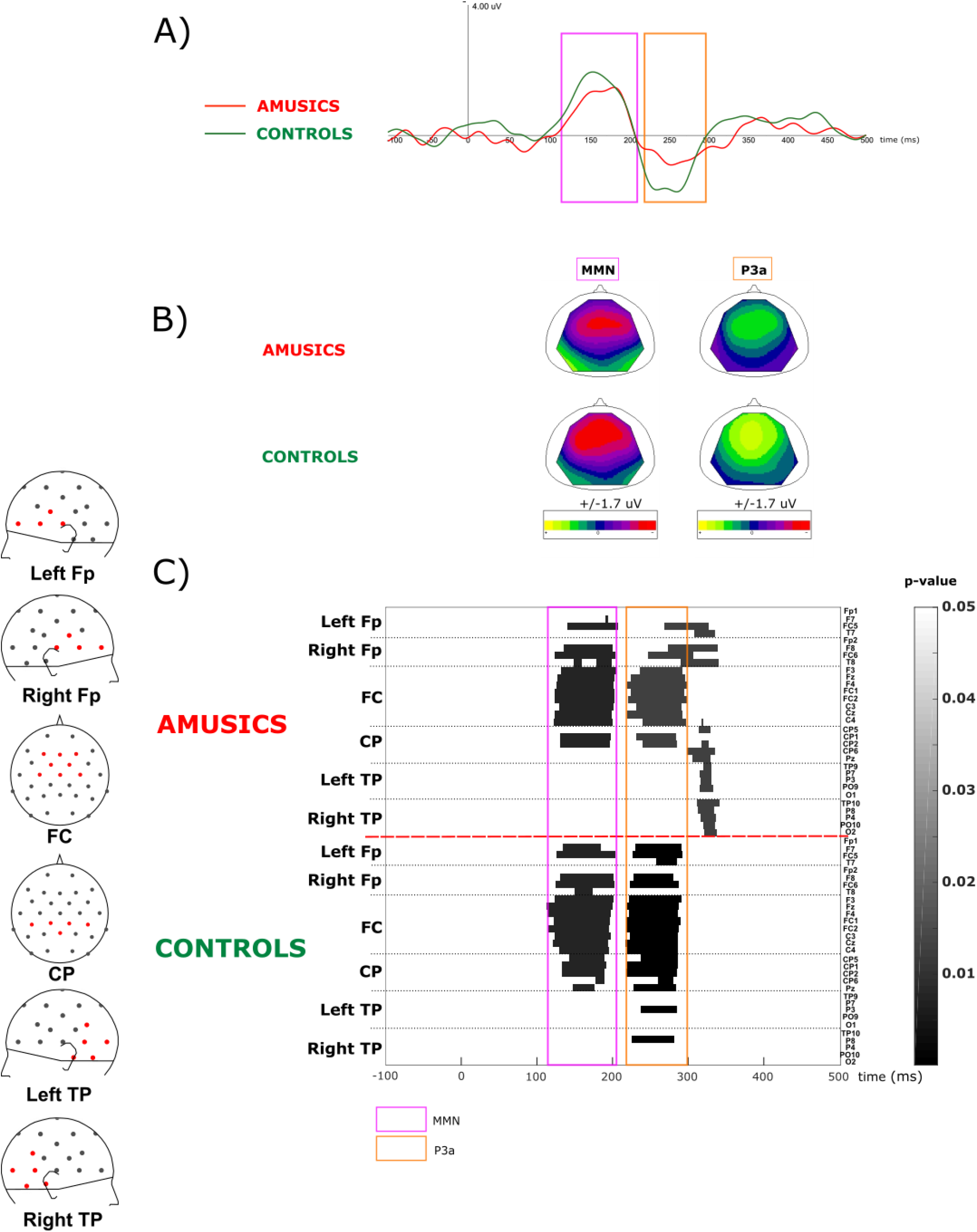
Evoked response to an emotional anger deviant in Amusics and Controls. **A)** Average curve of eight fronto-central electrodes (Fp1, Fp2, F3, F4, FC1, FC2, C3, C4) of the response to the anger deviant minus the response to the neutral standard, for amusics and controls, negativity is up. **B)** Topographies for the two evoked potentials (MMN and P3) over the emergence windows identified below, separately for amusics and controls. Amplitude scale is indicated for each ERP. **C)** Emergence of evoked responses for amusics and controls for each electrode, grouped by topography, emergence windows used for the analysis are in pink for the MMN (113-205ms), orange for the P3a (217-299ms). Fp= pre-frontal, FC=fronto-central, CP=centro-parietal, TP=temporo-parietal.

#### 2. MMN

After comparison to the null model, the best model showing decisive evidence was the model with the main effect of Localization (BF10=2.79e+10). This model was 1.64 times better than the model with the main effects of Localization and Group (BF10=1.7e+10), and 3.79 times better than the model with the main effects of Localization and Side (BF10=7.36e+9). The best model was at least 6.2 times better than the other models (BF10<4.52e+9). This was confirmed by a decisive specific effect of Localization (BFinclusion=1.2e+10), and no other specific effects (BFinclusion<0.29). The Group effect emerging in the second best model showed that the amusics tended to have a smaller MMN than controls. According to post-hoc tests, amplitudes at Fp and C were smaller than amplitudes at FC and F (all pcorr<0.004).

#### 3. P3a

After comparison to the null model, the best model showing decisive evidence was the model with the main effect of Localization (BF10=6.25e+15). This model was 1.63 times better than the model with the main effects of Localization and Group (BF10=3.83e+15), and 7.98 times better than the model with the main effects of Localization and Side (BF10=7.83e+14). The best model was at least 14 times better than the other models (BF10<4.44e+14). This was confirmed by a decisive specific effect of Localization (BFinclusion=∞), and no other specific effects (BFinclusion<0.23). The Group effect emerging in the second best model showed that amusics tended to have a smaller P3a than controls. According to post-hoc, amplitude at Fp was smaller than amplitudes at F, C and FC (all pcorr<0.015), amplitude at C was smaller than amplitudes at F and FC (both pcorr<0.041).

### Response to an anger deviant (Figure3)

#### 1. MMN

After comparison to the null model, the best model showing decisive evidence was the model with the main effect of Localization (BF10=5.99e+14). This model was 1.74 times better than the model with the main effects of Localization and Group (BF10=3.45e+14), and 4.68 times better than the model with the main effects of Localization and Group and the interaction between the two (BF10=1.28e+14). The best model was at least 8.14 times better than the other models (BF10<7.36e+14). This was confirmed by a decisive specific effect of Localization (BFinclusion=9.46e+13), and no other specific effects (BFinclusion<0.3). The Group effect emerging in the second best model showed that amusics tended to have a smaller MMN than controls. According to post-hoc, amplitude at Fp sites was smaller than amplitudes at C, FC and F (all pcorr<0.006), amplitude at C and F were smaller than amplitude at FC (both pcorr<0.035).

#### 2. P3a

After comparison to the null model, the best model showing decisive evidence was the model with the main effects of Localization and Group and the interaction between the two (BF10=3.25e+16). This model was 4.6 times better than the model with the main effects of Localization, Side, Group, and the interaction between Localization and Group (BF10=7.02e+15), and 6.1 times better than the model with the main effects of Localization and Group (BF10=5.3e+15). The best model was at least 11.9 times better than the other models (BF10<2.95e+15). This was confirmed by a decisive specific effect of Localization (BFinclusion=∞), and positive effects of Group (BFinclusion=4.99) and the interaction between Localization and Group (BFinclusion=9.23), and no other specific effects (BFinclusion<0.09). According to post-hoc tests, amplitude at Fp was smaller than amplitudes at C, FC and F (all pcorr<0.017), amplitudes at C and F were smaller than amplitude at FC (both pcorr<0.001), amplitude at C was smaller than amplitude at F (pcorr=0.032). Amusics had a significantly smaller P3 compared to Controls. This group difference was especially observed at Fp sites (pcorr=0.081). Specifically, amusics had smaller amplitude at Fp compared to amplitudes at F, FC, C (all pcorr<0.001), whereas controls had smaller amplitude at Fp compared only to FC (pcorr=0.002).

### Comparisons between deviants

To investigate potential differences across emotions, we ran a Bayesian ANOVA with the additional within-subjects factor Emotion for each evoked potential.

#### 1. Early negativity

This analysis included only the neutral and sadness deviants. After comparison to the null model, the best model showing decisive evidence was the model with the main effects of Localization and Emotion (BF10=3.2e+10). This model was 1.4 times better than the model with the main effects of Localization, Emotion, Group and the interaction between Localization and Group (BF10=2.3e+10), and 1.6 times better than the model with the main effects of Localization, Emotion and Group (BF10=2.05e+10). The best model was at least 4.1 times better than the other models (BF10<7.8e+9). This was confirmed by a decisive specific effect of Localization (BFinclusion=6.17e+8), a small positive effect of Emotion (BFinclusion=2.68) and no other specific effects (BFinclusion<0.51). According to post-hoc tests, the amplitude at Fp sites was smaller than amplitudes at F, FC and C (all pcorr<0.012), amplitude at FC was larger than amplitudes at F and C (all pcorr<0.001). The early negativity for sadness was smaller than the one for neutrality. Amusics tended to have a smaller early negativity than controls. Specifically, amusics tended to have smaller amplitude at Fp compared to amplitudes at F, FC, C (all pcorr<0.001), whereas controls tended to have smaller amplitudes at Fp compared to amplitudes at FC only (pcorr=0.019).

### 2. MMN

After comparison to the null model, the best model showing decisive evidence was the model with the main effects of Localization and Emotion (BF10=1.6e+18). This model was 2.2 times better than the model with the main effects of Localization, Emotion and Group (BF10=7.15e+17), and 13.3 times better than the model with the main effects of Localization, Emotion and Side (BF10=1.2e+17). The best model was at least 18.8 times better than the other models (BF10<8.49e+16). This was confirmed by a decisive specific effect of Localization (BFinclusion=2.82e+13) and Emotion (BFinclusion=782), and no other specific effects (BFinclusion<0.071). The Group effect emerging in the second best model showed that amusics tended to have a smaller MMN than controls. According to post-hoc tests, amplitude at Fp and C sites was smaller than amplitudes at F, FC (all pcorr<0.001), amplitude at Fp was smaller than amplitude at C (pcorr=0.029). The MMN tended to be smaller for sadness compared to anger (pcorr=0.2).

#### 3. P3a

After comparison to the null model, the best model showing decisive evidence was the model with the main effects of Localization, Emotion, Side, and Group, and the interaction between Emotion and Localization, between Emotion and Side, between Side and Localization, between Emotion and Group, and the triple interaction between Emotion, Localization and Side (BF10=9.85e+82). This model was 16.2 times better than the model with the main effects of Localization, Emotion, Side and Group and the interaction between Emotion and Localization, between Emotion and Side, between Side and Localization, between Emotion and Group, between Side and Group, and the triple interaction between Emotion, Localization and Side (BF10=6.08e+81), and 78 times better than the model with the main effects Localization, Emotion, Side and Group and the interaction between Emotion and Localization, between Emotion and Side, between Side and Localization, between Emotion and Group, between Localization and Group, and the triple interaction between Emotion, Localization and Side (BF10=1.26e+81). The best model was at least 99 times better than the other models (BF10<9.9e+80). This was confirmed by a decisive specific effect of Localization (BFinclusion=∞), Emotion (BFinclusion=∞), Side (BFinclusion=1164), Group (BFinclusion=6949), the interaction between Emotion and Localization (BFinclusion=1.24e+10), the interaction between Emotion and Side (BFinclusion=664), the interaction between Emotion and Group (BFinclusion=33214), the interaction between Localization and Side (BFinclusion=676) and the interaction between Emotion, Localization and Side (BFinclusion=4740), and no other specific effects (BFinclusion<0.07). Amusics had a smaller P3a compared to Controls. According to post-hoc tests, amplitude at Fp was smaller than amplitudes at C, FC and F (all pcorr<0.016), amplitudes at C and F were smaller than amplitude at FC (both pcorr<0.016), amplitude at C was smaller than amplitude at F (pcorr<0.001). The P3a was larger for anger compared to sadness and neutrality (both pcorr<0.046), and larger for sadness compared to neutrality (pcorr<0.001). More specifically, amplitudes for neutrality was smaller than amplitude for anger and sadness at C, F and FC (all pcorr<0.002), amplitude for neutrality was smaller than amplitude for anger at Fp (pcorr=0.002). Neutrality had smaller amplitude than had sadness and anger for both left and right sides (all pcorr<0.001). Amplitudes were larger over left side than right side. This was driven by a smaller amplitude at C for right side compared to left side (pcorr=0.098). This difference was driven in particular by the difference between the two groups for anger (pcorr=0.1). In amusics, neutrality had smaller amplitude than had anger (pcorr=0.025), whereas in controls, neutrality had smaller amplitude than had anger and sadness (both pcorr<0.004).

The analyses of components amplitude at midline electrodes (Fz, Cz, Pz) are reported in the supplementary material. Only limited group effects were observed in these analyses, in keeping with the results reported above which reveal that between-group differences were mostly observed at prefrontal sites, for which we did not have a midline electrode in our 32-electrode montage. These results at midline electrodes further emphasize that the early negativity peaking at Fz and Cz was slightly more central than the MMN, which peaked at Fz. This is in agreement with the hypothesis that the early negativity included N1 refractoriness effects.

## Discussion

Using an oddball paradigm with emotional prosody stimuli, we revealed the automatic brain responses of congenital amusic individuals compared to matched control participants for neutral and emotional verbal sounds. Based on previous behavioral and ERP results, we expected a decreased early automatic processing of deviancy in amusics compared to controls, with potentially different responses to neutral and emotional deviancy in these two groups. Amusics had reduced automatic processing of a neutral deviant compared to controls, with a diminished early negativity at the latency of N1 and a slightly reduced MMN. Similarly, the early processing of emotional stimuli (reflected by the early negativity at the latency of N1) was decreased in amusics compared to controls, yet with only slightly reduced emotional MMNs. The later P3a observed in response to a salient emotional deviant (anger) was strongly decreased in amusics compared to controls. These results suggest a differential processing of neutrality and emotions, with impaired pre-attentive processing of both neutral and emotional sounds in congenital amusia, at early cortical processing stages (around 100 ms) and in late processing stages associated with high-level cognitive processes (around 300 ms). The rather preserved MMN in between these altered processing stages suggest that change detection mechanisms can operate on degraded initial sound representations, at least in the case of large enough sound deviances.

Even if congenital amusia was first described to be music-specific (Ayotte et al., 2002; Peretz et al., 2003), recent evidence suggest that the pitch deficit in congenital amusia could also extend to speech material, even though to a lesser extent (Nguyen et al., 2009; Tillmann, Burnham, et al., 2011; Tillmann, Rusconi, et al., 2011; Zhang et al., 2017). In relation with the present study, congenital amusia is not only a music perception deficit but also a language processing deficit, in particular for non-verbal auditory cues such as emotional prosody (Lolli et al., 2015; Nguyen et al., 2009; Patel et al., 2008; Pralus et al., 2019; Thompson et al., 2012).

### Impaired early encoding of auditory stimuli in congenital amusia

A smaller early negativity was observed in amusics compared to controls for neutral and sadness deviants. It points to amusics’ increased difficulties to automatically process the deviants at early processing stages. This early negativity seems to correspond to N1 adaptation effects as the adaptation observed here occurs in the latency range of the N1, with a slightly different topography than the subsequent MMN.

In agreement with previous research (Albouy et al., 2013; Omigie et al., 2013), our results thus reveal an early deficit of auditory encoding in the amusics’ brain. This early processing seems to be particularly less efficient for neutral stimuli, but can also be altered for emotional stimuli, as revealed by the results with the sadness deviant. Interestingly, taken together, the results suggest a general decrease of the early negativity in congenital amusia, observed both in the processing of pitch sequences (Albouy et al., 2013; Omigie et al., 2013) and in oddball contexts (current results). As suggested by stimulus-specific adaptation research (Carbajal & Malmierca, 2018; Malmierca et al., 2014; Pérez-González & Malmierca, 2014), a precise representation of the standard is necessary to elicit a strong N1 when the deviant is presented. However, if the representation of the standard is not precise, as in congenital amusics, the N1 elicited by the deviant remains similar to the N1 elicited by the standard, as revealed here using an oddball paradigm.

Interestingly, similar pitch deviance was used with the three types of deviant (the smaller pitch deviance was for the sadness deviancy). Even though we tried to match the acoustic differences between the three types of deviant as closely as possible, other acoustic features than pitch differentiated between the three emotions. These variations of acoustic parameters could explain at least in part the pattern of evoked responses in the two groups. As roughness and inharmonicity were higher for the anger deviant compared to sadness and neutrality, it could have helped amusics to correctly process this anger deviant and recognize it behaviorally (Pralus et al., 2019). Indeed, previous reports suggest that amusics’ emotional judgments are based largely on roughness and tempo rather than harmonicity cues, which are mostly used by controls (Gosselin et al., 2015; Lévêque et al., 2018; Marin et al., 2015). Moreover, the anger deviant was characterized by a longer attack time, in particular when compared to the neutral standard. This could explain why the pattern of the first evoked potentials in response to this deviant was different compared to the two other emotions, with no early negativity at the latency of N1. However, this specific pattern of responses was similar in the two groups.

### Preserved automatic change detection and implicit processes in amusia

To investigate automatic change detection processes in congenital amusia, we studied the MMN evoked by the three deviants. As expected (Carminati et al., 2018; Charpentier et al., 2018), a MMN was induced by both emotional and neutral deviants, compared to a neutral standard in both groups. The MMN was larger for emotional deviants than for the neutral deviant (Schirmer et al., 2005, 2016).

No clear deficit of the MMN for the neutral or emotional deviants was observed in amusics compared to controls, suggesting at least a partially preserved automatic processing of emotional prosody in amusics, as previously shown with behavioral data (Lima et al., 2016; Lolli et al., 2015; Pralus et al., 2019; Thompson et al., 2012). This result is in line with previous research on automatic pitch processing in congenital amusia demonstrating only a small deficit of the congenital amusics’ MMN for small pitch changes in tone sequences (Fakche et al., 2018; Moreau et al., 2009, 2013; Nan et al., 2016; Omigie et al., 2013; Zhang & Shao, 2018). It suggests that, despite an impaired early processing of the deviant, congenital amusics’ brain is still able to automatically detect the change. However, even if it was not significant, we did observe a small decrease of the MMN to the neutral deviant in amusics, suggesting that this implicit knowledge in amusics’ brain might not be fully sufficient in some cases to allow the change detection mechanisms underlying the MMN to produce as large error signals as in controls. It is widely admitted that a correct sensory memory representation of the standard is needed to elicit an MMN (Näätänen et al., 2005). This would suggest that in congenital amusic participants, this memory representation is not as accurate as in controls. Thus, these results would contribute to the understanding of the deficit in congenital amusia as previously demonstrated with short-term memory tasks (Albouy et al., 2013, 2016; Fakche et al., 2018; Graves et al., 2019; Tillmann et al., 2009; Williamson & Stewart, 2010).

In combination with the analysis of the early negativity, these results show that acoustic sensitivity is impaired in congenital amusia, and do not seem to depend on emotional content of the stimulus. However, the more cognitive and memory-related comparison reflected by the MMN (Maess et al., 2007) seems to be less impaired in congenital amusia. In particular, it appears that this component would be only minimally impacted in congenital amusia when an emotional component is present in the stimulus. Such preserved automatic cortical processing steps could be the basis of the preserved implicit processes observed behaviorally in musical and emotional judgements (Lévêque et al., 2018; Pfeuty & Peretz, 2010; Pralus et al., 2019; Stewart, 2011; Tillmann et al., 2007; Tillmann, Lalitte, et al., 2016).

### Decreased awareness of emotional stimuli in congenital amusia

To further investigate the potential deficit of awareness in congenital amusia for emotional stimuli, we analyzed the P3a in response to the three types of deviants. This ERP was larger for the emotional deviants, especially for anger, but was still detectable for the two deviants in the two participant groups. For the anger deviant, which elicited the largest P3a in controls, the P3a was strongly decreased in amusics compared to controls. A reduced P3a in amusics was previously shown with lexical tones (Zhang & Shao, 2018) and using tasks with small pitch changes in tone sequences (Braun et al., 2008; Moreau et al., 2009, 2013). The decreased P3a relates to an awareness deficit suggested in congenital amusia (Peretz et al., 2009), in particular for emotional stimuli (Lévêque et al., 2018; Pralus et al., 2019). Specifically, P3a is considered to reflect automatic attentional orientation toward a salient deviant (Escera et al., 1998; Polich & Criado, 2006). Thus, congenital amusics would have a deficit to process unexpected novel sounds. However, amusics were still able to perform the recognition task for the anger deviant. These results suggest that when the automatic preattentional processes of the amusics reach a sufficient level (a sizeable MMN and a detectable P3a), they can perform the recognition task, despite this deficit at these late processing stages.

### Brain networks involved in emotional prosody perception in congenital amusia

The group differences were mostly visible on bilateral pre-frontal electrodes. Interestingly, in congenital amusia, frontal regions were found to be altered (Albouy et al., 2013, 2019; Hyde et al., 2006, 2007, 2011). In particular, decreased gray and white matter volume of the inferior frontal cortices was observed in congenital amusia (Albouy et al., 2013, 2019; Hyde et al., 2006, 2007, 2011). As these regions are involved in emotional prosody processing (Frühholz et al., 2012; Liu et al., 2015), it could have been expected that amusics would have a deficit to perceive emotional prosody. However, our results with the MMN suggest a partial preservation of these circuits to automatically detect emotional prosody in congenital amusics. These results are in line with previous reports showing that the perception of emotional prosody does not only involve a fronto-temporal network, but also extend to other regions, such as probably the amygdala that detects salience and meaningful information (Frühholz et al., 2016), which would be preserved in congenital amusia. Further research using brain imaging (with fMRI for example) should investigate the brain networks involved in emotional perception in congenital amusia.

### Conclusion

Our present findings shed new light on different aspects of automatic sound processing in congenital amusia, in particular for speech material and its emotional features. The observed impairments might lead to difficulties to process speech correctly in some situations. For instance, in degraded conditions such as hearing in noise, challenging conditions for speech comprehension (Liu et al., 2015; Oxenham, 2008, 2012; Tang et al., 2018), amusics could have more difficulties to understand the speaker’s emotions and intentions (Mcdonald & Stewart, 2008; Omigie et al., 2012). Moreover, this study gives further insight about the dissociation of implicit and explicit processing in congenital amusia (Lévêque et al., 2018; Omigie et al., 2013; Pralus et al., 2019; Tillmann et al., 2012, 2014; Tillmann, Lalitte, et al., 2016). It reveals the overall pattern of emotional perception in congenital amusia, from the first steps of cortical processing (Albouy et al., 2013; Omigie et al., 2013) to the late processing stages (P300, Moreau et al., 2009, 2013; Peretz et al., 2009), via the intermediate stage of change detection reflected by the MMN. This relatively preserved MMN might relate to preserved implicit processing in congenital amusia for music and emotional prosody stimuli.

## Fundings

This work was conducted in the framework of the LabEx CeLyA (‘‘Centre Lyonnais d’Acoustique”, ANR-10-LABX-0060) and of the LabEx Cortex (‘‘Construction, Cognitive Function and Rehabilitation of the Cortex”, ANR-11-LABX-0042) of Université de Lyon, within the program ‘‘Investissements d’avenir” (ANR-16-IDEX-0005) operated by the French National Research Agency (ANR).

## Footnotes

1 The equiprobable stimuli could not be used as the reference stimulus to compute difference ERPs, as in the equiprobable block, the anger sound (and to a lesser extent the neutral deviant sound) elicited a negativity compared to the other equiprobable sounds in the latency range of the MMN, suggesting that MMNs were elicited within this sequence (see Figure S1).

2 We also computed Bayesian ANOVA on the amplitude at midline electrodes (Fz, Cz, Pz) for each time-window for each emotion. See Supplementary analysis for details.

## Notes

### Competing Interest Statement

The authors have declared no competing interest.

