## Supplementary for "Pre-attentive processing of neutral and emotional sounds in congenital amusia"

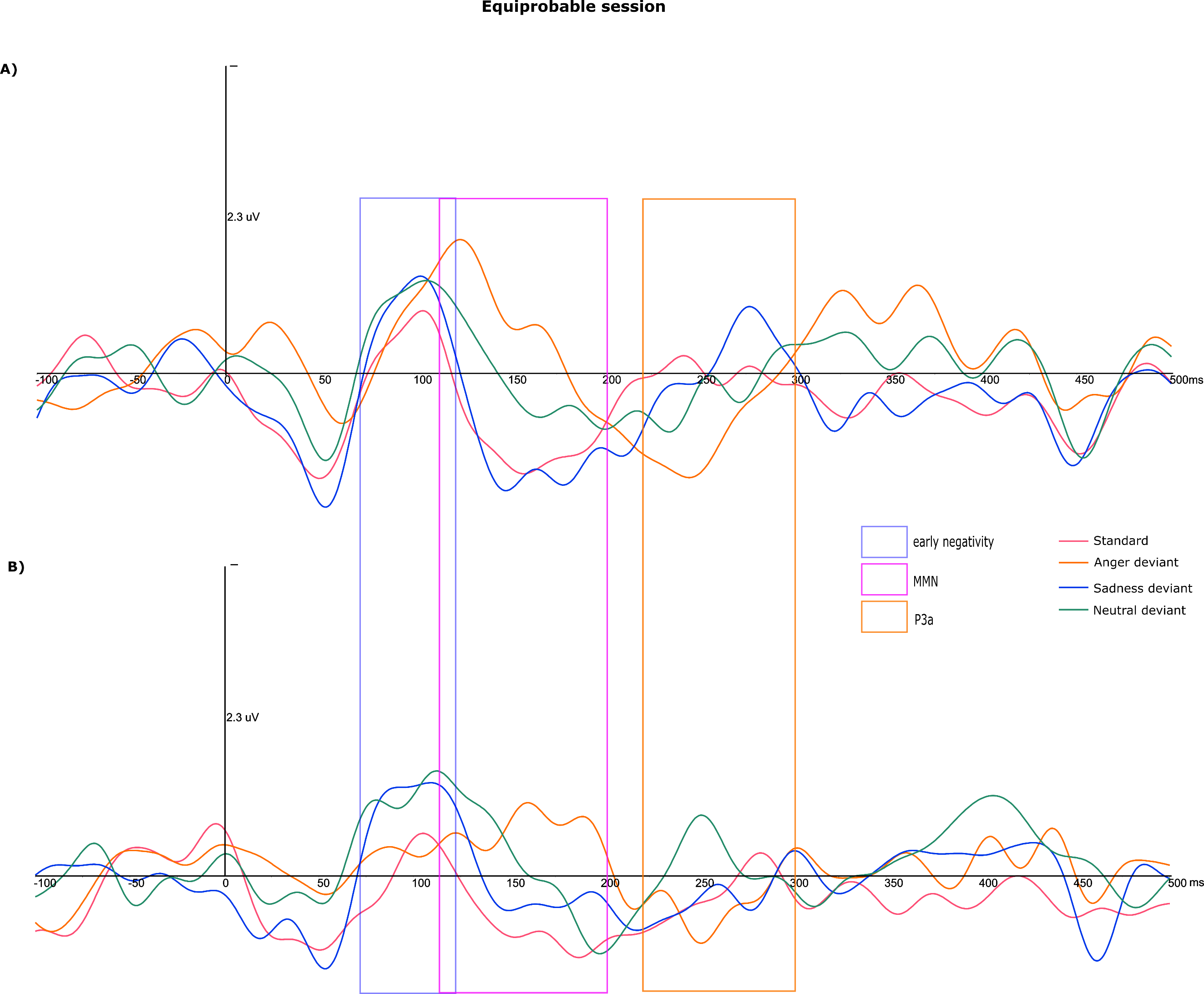

**Figure S1:** Average curve of ten fronto-central electrodes (Fp1, Fp2, F3, Fz, F4, FC1, FC2, C3, Cz, C4) of the response in the equiprobable session to the sounds that were used as a neutral standard, anger, sadness and neutral deviants in the main blocks, for controls (A) and amusics (B). Negativity is up. Emergence windows of the principal analysis are reported. The anger sound (and to a lesser extent the neutral deviant sound) elicited a negativity compared to the other equiprobable sounds in the latency range of the MMN, suggesting that MMNs were elicited within this sequence.

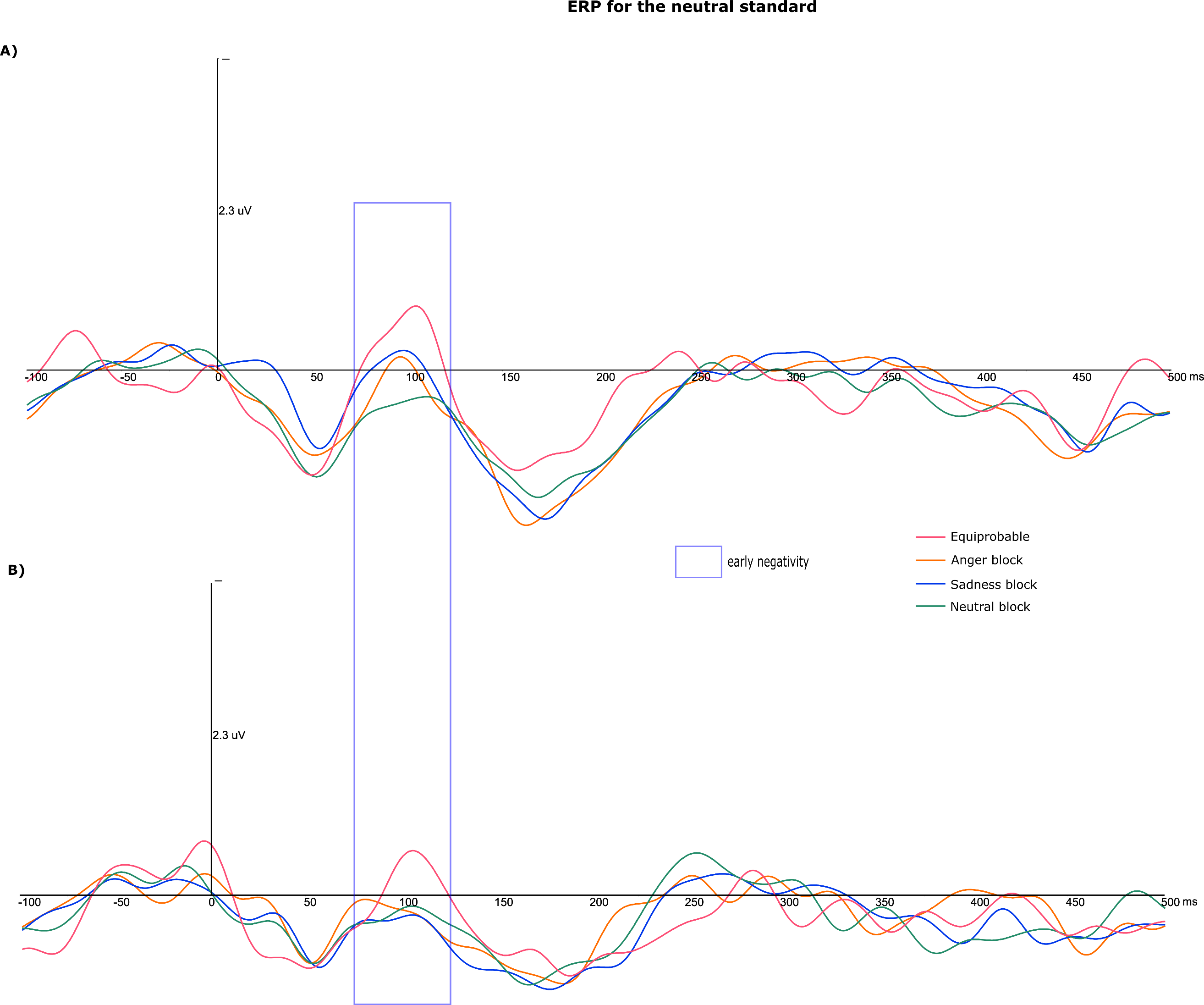

**Figure S2:** Average curve of ten fronto-central electrodes (Fp1, Fp2, F3, Fz, F4, FC1, FC2, C3, Cz, C4) of the response to a neutral standard in the four conditions (equiprobable session, block 1 with anger deviant, block 2 with sadness deviant, block 3 with neutral deviant), for controls (A) and amusics (B). Negativity is up. Emergence window or the early negativity in the principal analysis is reported. An early negativity at the latency of N1 was present only in the equiprobable session, suggesting that a habituation of N1 was occurring in the oddball blocks.

### Supplementary analysis on midline electrodes (Fz, Cz, Pz)

#### Response to a neutral deviant

##### 1. Early negativity

After comparison to the null model, the best model showing decisive evidence was the model with the main effect of Localization (BF10=1.3e+5). This model was 1.9 times better than the model with the main effects of Localization and Group (BF10=6.7e+4), and 8.7 times better than the model with the main effects of Localization and Group and the interaction between the two (BF10=1.5+4). The model with the main effect of Group showed no evidence (BF10=0.45). This was confirmed by a decisive specific effect of Localization (BFinclusion=9.6e+4), and no other specific effects (BFinclusion<0.43). The Group effect emerging in the second best model showed that amusics tended to have a smaller early negativity than controls. According to post-hoc tests, amplitude at Pz was smaller than amplitudes at Fz and Cz (both pcorr<0.001).

##### 2. MMN

After comparison to the null model, the best model showing decisive evidence was the model with the main effect of Localization (BF10=1.7e+11). This model was 1.6 times better than the model with the main effects of Localization and Group (BF10=1.04e+11), and 1.7 times better than the model with the main effects of Localization and Group and the interaction between the two (BF10=1+11). The model with the main effect of Group showed no evidence (BF10=0.51). This was confirmed by a decisive specific effect of Localization (BFinclusion=1.7e+11), and no other specific effects (BFinclusion<1.4). The Group effect emerging in the second best model showed that amusics tended to have a smaller MMN than controls. According to post-hoc tests, amplitude at Pz was smaller than amplitudes at Fz and Cz (both pcorr<0.001), amplitude at Cz was smaller than amplitude at Fz (pcorr<0.001).

### 3. P3a

After comparison to the null model, the best model showing decisive evidence was the model with the main effect of Localization (BF10=4.1e+8). This model was 2.4 times better than the model with the main effects of Localization and Group (BF10=1.7e+8), and 15.8 times better than the model with the main effects of Localization and Group and the interaction between the two (BF10=2.6e+7). The model with the main effect of Group showed no evidence (BF10=0.36). This was confirmed by a decisive specific effect of Localization (BFinclusion=2.9e+8), and no other specific effects (BFinclusion<0.3). The Group effect emerging in the second best model showed that amusics tended to have a smaller P3a than controls. According to post-hoc tests, amplitude at Pz was smaller than amplitudes at Fz and Cz (both pcorr<0.001), amplitude at Cz was smaller than amplitude at Fz (pcorr<0.001).

#### Response to a sadness deviant

##### 1. Early negativity

After comparison to the null model, the best model showing decisive evidence was the model with the main effect of Localization (BF10=2.1+6). This model was 1.5 times better than the model with the main effects of Localization and Group (BF10=1.4e+6), and 4.1 times better than the model with the main effects of Localization and Group and the interaction between the two (BF10=5.1e+5). The model with the main effect of Group showed no evidence (BF10=0.62). This was confirmed by a decisive specific effect of Localization (BFinclusion=1.7e+6), and no other specific effects (BFinclusion<0.6). The Group effect emerging in the second best model showed that amusics tended to have a smaller early negativity than controls. According to post-hoc tests, amplitude at Pz was smaller than amplitudes at Fz and Cz (both pcorr<0.001).

##### 2. MMN

After comparison to the null model, the best model showing decisive evidence was the model with the main effect of Localization (BF10=3.9e+9). This model was 2.1 times better than the model with the main effects of Localization and Group (BF10=1.9e+9), and 9.7 times better than the model with the main effects of Localization and Group and the interaction between the two (BF10=4.02+8). The model with the main effect of Group showed no evidence (BF10=0.43). This was confirmed by a decisive specific effect of Localization (BFinclusion=2.9e+9), and no other specific effects (BFinclusion<0.4). The Group effect emerging in the second best model showed that amusics tended to have a smaller MMN than controls. According to post-hoc tests, amplitude at Pz was smaller than amplitudes at Fz and Cz (both pcorr<0.001), amplitude at Cz was smaller than amplitude at Fz (pcorr<0.001).

### 3. P3a

After comparison to the null model, the best model showing decisive evidence was the model with the main effect of Localization (BF10=6.2e+6). This model was 1.7 times better than the model with the main effects of Localization and Group (BF10=3.6e+6), and 12.1 times better than the model with the main effects of Localization and Group and the interaction between the two (BF10=5.1e+5). The model with the main effect of Group showed no evidence (BF10=0.49). This was confirmed by a decisive specific effect of Localization (BFinclusion=4.6e+6), and no other specific effects (BFinclusion<0.4). The Group effect emerging in the second best model showed that amusics tended to have a smaller P3a than controls. According to post-hoc tests, amplitude at Pz was smaller than amplitudes at Fz and Cz (both pcorr<0.001), amplitude at Cz was smaller than amplitude at Fz (pcorr=0.012).

#### Response to an anger deviant

##### 1. MMN

After comparison to the null model, the best model showing decisive evidence was the model with the main effect of Localization (BF10=8.5e+10). This model was 1.7 times better than the model with the main effects of Localization and Group (BF10=4.9e+10), and 11.6 times better than the model with the main effects of Localization and Group and the interaction between the two (BF10=7.3+9). The model with the main effect of Group showed no evidence (BF10=0.48). This was confirmed by a decisive specific effect of Localization (BFinclusion=6.4e+10), and no other specific effects (BFinclusion<0.4). The Group effect emerging in the second best model showed that amusics tended to have a smaller MMN than controls. According to post-hoc tests, amplitude at Pz was smaller than amplitudes at Fz and Cz (both pcorr<0.001), amplitude at Cz was smaller than amplitude at Fz (pcorr=0.016).

### 2. P3a

After comparison to the null model, the best model showing decisive evidence was the model with the main effects of Localization and Group (BF10=1.8e+10). This model was 1.2 times better than the model with the main effect of Localization (BF10=1.5e+10), and 7.8 times better than the model with the main effects of Localization and Group and the interaction between the two (BF10=2.3e+9). The model with the main effect of Group showed no evidence (BF10=1.3). This was confirmed by a decisive specific effect of Localization (BFinclusion=1.02e+10), and no other specific effects (BFinclusion<0.9). The Group effect emerging in the second best model showed that amusics had a smaller P3a than controls. According to post-hoc tests, amplitude at Pz was smaller than amplitudes at Fz and Cz (both pcorr<0.001).
